# Improved accuracy of lesion to symptom mapping with multivariate sparse canonical correlations

**DOI:** 10.1101/149260

**Authors:** Dorian Pustina, Brian Avants, Olufunsho Faseyitan, John Medaglia, H. Branch Coslett

## Abstract

Lesion to symptom mapping (LSM) is a crucial tool for understanding the causality of brain-behavior relationships. The analyses are typically performed by applying statistical methods on individual brain voxels (VLSM), a method called the mass-univariate approach. Several authors have shown that VLSM suffers from limitations that may decrease the accuracy and reliability of the findings, and have proposed the use of multivariate methods to overcome these limitations. In this study, we propose a multivariate optimization technique known as sparse canonical correlation analysis for neuroimaging (SCCAN) for lesion to symptom mapping. To validate the method and compare it with mass-univariate results, we used data from 131 patients with chronic stroke lesions in the territory of the middle cerebral artery, and created synthetic behavioral scores based on the lesion load of 93 brain regions (putative functional units). LSM analyses were performed with univariate VLSM or SCCAN, and the accuracy of the two methods was compared in terms of both overlap and and displacement from the simulated functional areas. Overall, SCCAN produced more accurate results - higher dice overlap and smaller average displacement - compared to VLSM. This advantage persisted at different sample sizes (N=20-131) and different multiple comparison corrections (false discovery rate, FDR; Bonferroni; permutation-based family wise error rate, FWER). These findings were replicated with a fully automated SCCAN routine that relied on cross-validated predictive accuracy to find the optimal sparseness value. Simulations of one, two, and three brain regions showed a systematic advantage of SCCAN over VLSM; under no circumstance could VLSM exceed the accuracy obtained with SCCAN. When considering functional units composed of multiple brain areas VLSM identified fewer areas than SCCAN. The investigation of real scores of aphasia severity (aphasia quotient and picture naming) showed that SCCAN could accurately identify known language-critical areas, while VLSM either produced diffuse maps (FDR correction) or few scattered voxels (FWER correction). Overall, this study shows that a multivariate method, such as, SCCAN, outperforms VLSM in a number of scenarios, including functional dependency on single or multiple areas, different sample sizes, different multi-area combinations, and different thresholding mechanisms (FWER, Bonferroni, FDR). These results support previous claims that multivariate methods are in general more accurate than mass-univariate approaches, and should be preferred over traditional VLSM approaches. All the methods described in this study are available in the newly developed LESYMAP package for R.

## INTRODUCTION

Lesion to symptom mapping (LSM) is the primary tool researchers use to make inferences on the causality of brain relationships with behavior. Unlike correlations between behavior and locus of activation in functional neuroimaging, the presence of a brain lesion provides strong evidence of the critical involvement of a brain region on the cognitive deficit it causes. Modern brain stimulation techniques such as transcranial magnetic stimulation produce transient alterations of brain function that may also demonstrate a causal relationship between brain anatomy and behavior. These studies, however, are limited by the restricted extent of the effects (e.g., approximately 1 cubic cm of tissue) and the limited understanding of the effects of non-invasive brain stimulations (Farah, 2015; Farah et al., 2014; R. Hamilton et al., 2011). Thus, LSM currently remains a crucial and prevalent method for studying the causality of brain-behavior relationships.

Since the dawn of lesion studies, LSM methods have focused on studying groups of patients (i.e., Broca presented three different cases in his 1861 paper) (Broca, 1861). In modern times, group LSM studies were initially performed by overlapping lesions of patients with similar deficits (Blunk et al., 1981; Dronkers, 1996), and later by applying statistical methods on individual brain voxels (Bates et al., 2003; Rorden et al., 2007). The separate analysis of each voxel is called “the massively univariate (or mass-univariate) approach”. In recent years, it has become evident that the mass-univariate approach suffers from limitations that can severely compromise the ability to pinpoint cognitive functions to specific brain areas (Karnath and Smith, 2014; Kimberg et al., 2007; Mah et al., 2014; Nachev, 2015; Sperber and Karnath, 2017). Some of the limiting factors are beyond the researcher’s control and cannot be easily eliminated; e.g., the fact that neighboring voxels are frequently correlated with each other, or the non-random nature of lesions that follow the vascular anatomy. In our experience, the correlation between immediately neighboring voxels can be around r = 0.88 (computed from a dataset of 131 patients, see Supplementary Material: Neighborhood Correlation Maps). Other limitations emerge because the methods used for analyses are inappropriate. For example, the mass-univariate approach performs many thousands of tests, increasing dramatically the potential for false positives. This “noise” cannot be fully eliminated without removing some of the true signal (Bennett et al., 2009). The very term “voxel-based” is misleading in a strict sense, because neighboring voxels often have identical patterns, and produce the same exact statistic. A group of voxels with identical lesions in every subject constitute a “unique patch” (Kimberg et al., 2007; Rorden et al., 2009). A quick computation on the data utilized in this study shows that the real number of unique single voxels in a sample 131 patients is less than half the total number of analyzed voxels, and can be only worse with fewer subjects (see Methods). Recognizing the lack of distinct voxel-wise information, most software analyze unique patches instead of single voxels (e.g., Mricron and voxbo packages, (Kimberg et al., 2007; Rorden et al., 2007)). Another problem of the mass-univariate approach is that within a group of subjects different voxels have different lesion ratios, and, consequently, different statistical power. Effects located in high power areas are more likely to be detected, and might ultimately bias the overall statistical maps towards these areas (Kimberg et al., 2007; Mah et al., 2014). Currently, there is no solution to correct for the variation in power. The resulting displacement that may occur can lead to irreproducible results, or, even worse major interpretation errors (Mah et al., 2014; Nachev, 2015; Sperber and Karnath, 2017).

Several authors have recently suggested that multivariate methods may overcome the above limitations. The switch to multivariate analyses is not a mere switch in statistical software, but a choice motivated by important considerations. To understand this, a distinction must be made between measurement units and functional units. A measurement unit within the imaging space is a voxel, which can be 1mm, 2mm, or perhaps 0.2mm. The smallest unit of interest that can be measured in the brain is a single neuron. The goal of cognitive neuroscientists is to identify functional units, or areas of the brain that perform specific cognitive processes. Importantly, measurement units cannot be treated as functional units without falling in conceptual fallacies. The death of a single neuron is unlikely to produce hemiparesis. The lesion of a single voxel is unlikely to produce aphasia. Behavioral deficits emerge because a neuron or a voxel is affected within a broader context of many other lesioned neurons or voxels. In practical terms, this means that the identification of the functional area cannot be achieved unless measurement units are considered together as a whole. Mass-univariate analyses consider measurement units as individual functional units out of their context. A real world example would be like trying to identify city boundaries by mapping the presence of cars every 100 yards across the whole country. Inevitably this approach is less precise than the consideration of cars within a context (i.e., in a city many cars are found next to each other). Multivariate methods more directly model functional units as composites of measurement units, and can understand that the status of several measurement units, taken together, is important to determine a behavioral deficit. This concept is not limited to functional units composed of adjacent voxels, but also to units composed of voxels located far from each other (i.e., two areas of the brain acting as a single functional unit). Importantly, two brain regions can interact towards the same goal in different ways (Godefroy et al., 1998), i.e., as serial processing units (i.e., phonologic, lexical, or semantic retrieval in a picture naming task, (Dell et al., 1997; Schwartz et al., 2006/2)), or as parallel processing units (i.e., dorsal and ventral language pathways, (Hickok and Poeppel, 2004; Schwartz et al., 2006/2)). For this reason, attempts to model multi-area functional units should also consider several combination rules.

In this study, we propose the use of a multivariate LSM approach based on sparse canonical correlation analysis for neuroimaging (SCCAN). Canonical correlations take two sets of variables (i.e., behavioral scores and voxel values) and search for optimal weights to apply on each side such that one or more components are created that correlate with each other (Hotelling, 1936; Kuhn and Johnson, 2013; Stevens, 2009). In a simplified view, these components can be thought of as principal components (PCA) of the covariance matrix computed between two different modalities acquired in the same subjects. SCCAN represents an extension of canonical correlations analysis with additional constraints that are specific to neuroimaging data (Avants et al., 2014, 2010). In this respect, SCCAN combines (optionally) spatial smoothing, cluster thresholding, and positivity constraint to improve the interpretability of the components and prevent overfitting in the *p >> n* (more variables than subjects) context of most neuroimaging studies. From a broad perspective, SCCAN is considered an optimization algorithm (projected gradient descent) that finds a locally optimal solution to a regularized variant of the objective function defined by Hotelling in 1936 (Hotelling, 1936). The advantage of this approach, which all multivariate methods have in common, is the consideration of all measurement units together in a single model.

The main aim of this paper was to test whether a multivariate method, such as SCCAN, can outperform univariate analyses, as suggested by previous publications (Mah et al., 2014; Sperber and Karnath, 2017; Zhang et al., 2014). This aim is posed as a series of questions that compare the methods at different sample sizes, different thresholding mechanisms, and different hypothetical functional interactions between brain regions. Most of the comparisons were performed on results obtained from simulated brain-behavior relationships, that is, by using the real lesion maps of patients with stroke but synthetic behavioral scores that followed the lesion load of specific brain areas (putative functional units). We then tested the accuracy to which the source of the behavioral deficit could be identified with the different methods. The approach is similar to previous simulations studies (Mah et al., 2014; Sperber and Karnath, 2017; Zhang et al., 2014), though we used real functional areas derived from multiple atlases, and injected error in synthetic behavioral scores to match the notoriously noisy brain-behavior relationships.

A new R package was created during the course of this study, called LESYMAP. LESYMAP relies on ANTsR (Avants, 2015) for image manipulation, and includes all the necessary tools to perform real or simulated LSM analyses.

## METHODS

### Patients

A dataset of 131 lesion maps from patients with chronic stroke were included in the study (patient age 58.2 +/- 11.3, months post-onset 44 +/- 63, range 2 - 387 months; the “+/-” sign indicates standard deviation). This dataset is part of the Neuro-Cognitive Rehabilitation Research Patient Registry at the Moss Rehabilitation Research Institute, and has been previously used in other studies on post-stroke aphasia (Mirman et al., 2015; Pustina et al., 2016; Schwartz et al., 2012, 2011, 2009; Zhang et al., 2014). All lesions were located in the left hemisphere around the territory of the middle cerebral artery; no patients were included with stroke in other vascular territories. Lesion maps were derived from MRI (76 patients) or CT (55 patients) scans. MRI scans were acquired on a 3-T Siemens Trio scanner [repetition time (TR) = 1,620 ms, echo time (TE) = 3.87 ms, field of view (FOV) = 192 × 256 mm, 1 × 1 × 1-mm voxels]. CT scans were acquired using 60 axial 3 mm slices without using contrast agents. For patients with available MRI, lesions were manually drawn on the patient’s MRI, and then transferred in ch2 MNI template space using a series of linear and non-linear registrations in ANTs (Avants et al., 2011).

The lesioned area was removed from registration computations. For patients with CT scans, lesions were manually drawn on the ch2 MNI template by labeling the corresponding areas that appeared lesioned in the CT image. An expert neurologist with 30 years of experience (HBC) either manually drew or inspected the lesions drawn by people trained by him (all CT lesions were drawn by HBC). The average lesion size in template space was 100 +/- 82 ml (range: 5 - 376 ml).

Although we simulated the behavioral data, we used two real behavioral scores to estimate the amount of error to inject in brain-behavior relationships. The scores were (i) the aphasia quotient from the Western Aphasia Battery (WAB-AQ, (Kertesz, 1982)), and (ii) the number of correctly named items from Philadelphia Naming Test (PNTcorrect, (Roach et al., 1996)). The correlation between the two behavioral scores was r = 0.87. PNTcorrect had one missing data point, and the corresponding subject was excluded when analyzing this score.

### Patch-based descriptive statistics

A question we wanted to answer in this study was whether it is worth running LSM analyses on high resolution lesion maps. For example, the number of unique patches (i.e., groups of voxels with identical lesion pattern) might be similar at low and high resolution maps, causing unnecessary computational burden when using high resolution maps. To answer this question, we computed the number of unique patches (groups of voxels with the same identical lesion pattern) at 1mm, 2mm, and 3mm resolutions, at various sample sizes from N=20 to N=131 in steps of N=10. At each step we randomly sampled a group of subjects for 200 times and computed the average number of unique lesion patches. The last step contains all the subjects and was sampled only once.

### Defining functional areas

In order to achieve realistic simulations, the functional areas that generate the signal must be as close as possible in shape and size to real functional areas known from the literature. For this reason, we constructed a parcellation atlas by merging three publicly available atlases. The schematic pipeline is displayed in Figure 1. To parcellate cortical regions we used the Multi-Modal Parcellation (MMP) atlas obtained from Glasser et al. (2016) using data from Human Connectome Project. The authors derived this atlas using a robust estimation of functional areas from multiple imaging modalities, and validated the presence of the functional areas in new subjects through cross-validation procedures. A publicly available volumetric version of this atlas (http://neurovault.org/collections/1549/) was transformed from ICBM 2009a MNI space to ch2 MNI space using ANTs registration pipelines. The volumetric MMP atlas contains thin cortical parcels which might not overlap well with lesions due to registration errors of each subject. To give more consistency to the parcels, we extended each parcel to include small portions of the subcortical white matter immediately under the parcel. This was achieved with the `PropagateLabelsThroughMask` algorithm in ANTsR (radius=4). Subcortical nuclei were added from a second functional atlas, the Brainnetome atlas, which is also defined from multiple neuroimaging modalities (Fan et al., 2016). As a rule, only voxels left empty by the previous atlas were filled with the new parcels. Finally, white matter parcels were added by using the JHU atlas provided with FSL software (tract probability > 0.25, (Hua et al., 2008)).

**Figure 1.**
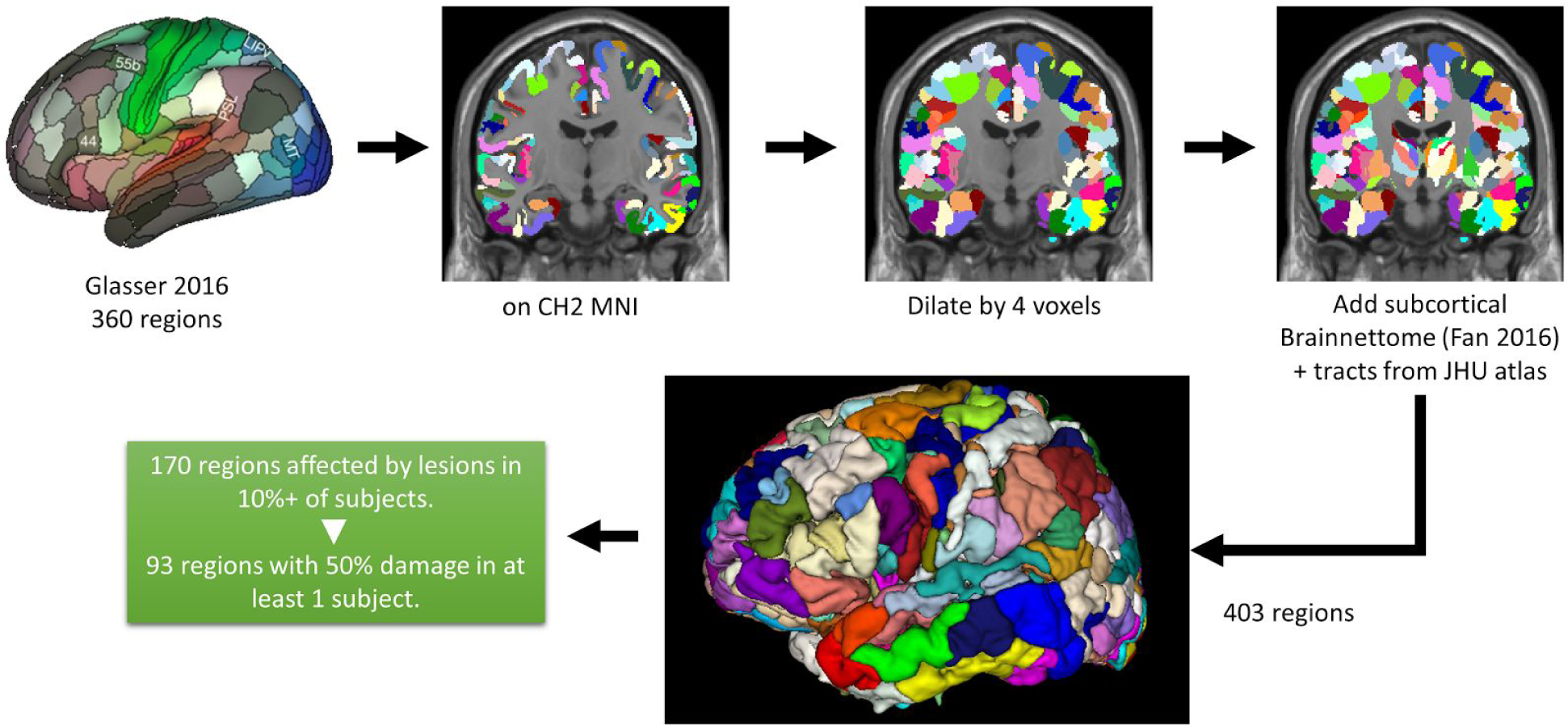
Schematic pipeline used to define functional areas.

After merging all three atlases, a total 403 different regions were defined across the brain. From this pool, we selected areas affected at least in one voxel in at least 10% of the patients, leaving a total of 170 regions. Preliminary analyses showed that many of these parcels were located at the margins of the vascular territory affected in our subjects, and were marginally affected in all subjects. This caused poor signal-to-noise ratio when error was injected into simulations, causing univariate analysis to fail. To avoid this issue, we applied a second inclusion rule to simulated parcels, such that only parcels affected 50% or more in at least one subject were included. The second rule reduced the total number of parcels to 93. The average volume of these 93 parcels was 2.8 +/- 1.8 ml. The lesion load of each parcel was computed as the ratio of lesioned voxels over the total number of voxels in the parcel (getLesionLoad in LESYMAP).

### Behavioral data simulation

A common aspect overlooked in simulation studies is the use of lesion load as a pure measure of behavioral deficit; for example, the more an area is lesioned the more a deficit arises. This perfect relationship is not realistic and can produce misleading simulations. From a statistical perspective, a perfect relationship creates large statistical scores that do not match the scores observed in real VLSM studies (i.e., peak t-score = 7 - 10). High statistical scores can affect corrections for multiple comparison, such as, false discovery rate (FDR, (Benjamini and Hochberg, 1995)). As a result, more extensive maps might be produced, and, consequently, the displacement of center of mass is overestimated. To achieve proper simulations we first estimated the typical peak t-scores from available behavioral scores: WAB-AQ and PNTcorrect. These two LSM analyses produced peak t-scores of 7.7 and 6.2, respectively, which are in the range of typical peak scores reported in the literature. With these values in mind, we aimed to simulate behavioral scores that would produce peak t-scores around these values. The formula for simulating the behavioral scores was “behavior = a*lesionLoad + b*error”, where a and b are weights. The sum of the weights is always 1, i.e., if a=0.7 then b=0.3. To establish the necessary amount of noise we performed the following empirical search. First, behavioral scores were simulated for each brain region using different amounts of error weight from 0 to 0.7 in steps of 0.1 (error weights above 0.7 produced often null results). Then, we performed VLSM analyses for 170 regions at each error level. Note, we used all affected regions to increase the chances of finding the highest possible t-score at every voxel. After completing a total of 1360 VLSM analyses using t-tests, a composite map was created at each error level by selecting, for each voxel, the maximum t-score across the 170 maps derived from the simulations. The map of highest t-scores at each voxel was used to compute an average and standard deviation of the t-scores expected for that error level. The result of this empirical search was Table 1. From this table it can be noted that errors around 0.5 - 0.6 produce t-scores of 9.5 and 7.5, respectively. Since the number of subjects in our study is high and the t-scores depend on the sample, we chose the 0.5 error weight for the remaining simulations.

**Table 1.**
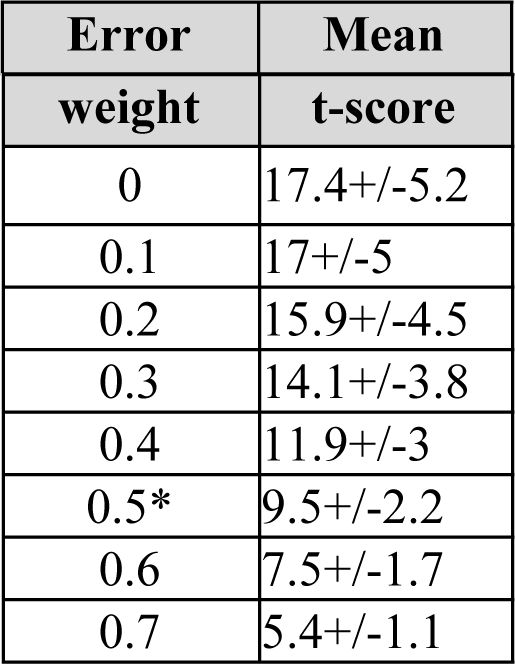
Average T-scores (+/- standard deviation) from simulations obtained at different error weights. The asterisk shows the chosen error weight for behavioral simulations.

### LSM analyses: mass-univariate

VLSM analyses are often performed using t-tests (Bates et al., 2003; Schwartz et al., 2012). However, these tests make specific assumptions that may not hold for all brain voxels (Rorden et al., 2007). A quick check of the assumptions of normality and variance homogeneity using our real data (WAB-AQ and PNTcorrect) showed that 13.4% of voxels violated the assumption of variance homogeneity, and 100% of voxels violated the assumption of the normality of distribution. Note, these assumptions depend on the distribution of behavioral scores, which can be difficult to predict for new samples or new cognitive batteries. To avoid checking for assumptions each time we performed a test on a voxel, we used the non-parametric Brunner-Munzel (BM) test (Brunner and Munzel, 2000). The BM test uses rank transformations of the data and is the method of choice in the MRIcron software (Rorden et al., 2007). Multiple comparison corrections were performed either with false discovery rate (Benjamini and Hochberg, 1995; Genovese et al., 2002), or with two different types of family wise error rate correction: Bonferroni (Bonferroni, 1936) or permutation based thresholding (Nichols and Holmes, 2002; Winkler et al., 2014), all thresholded at 0.05. In the remaining paragraphs, FWER will refer only to the permutation based correction, while Bonferroni will be called with its own name, despite both methods falling in the family of family wise error corrections. To compute the FWER threshold, we performed 1000 repetitions of the analyses with randomly permuted behavioral scores, and each time recorded the peak score (the voxel with maximal value). After creating a full distribution of the 1000 peak scores, the FWER threshold was set at the 95th percentile of the distribution. All univariate analyses were performed in LESYMAP.

### LSM analyses: multivariate

The sparseDecom2 function in ANTsR was used to run sparse canonical correlation analyses (SCCAN, (Avants et al., 2014, 2010)). SCCAN is an optimization procedure that gradually defines the weights to apply to each voxel such that the overall relationship with behavior is maximized while taking into account other constraints. Several regularization measures are integrated in SCCAN to factor spatially-clustered solutions over solutions with isolated voxels.

To achieve this, within the optimization algorithm, voxel weights are smoothed and clusters with just few voxels are set to zero by a user-defined regularization operator (a modified L1 penalty term). An important parameter in SCCAN is the sparseness value, which sets the proportion of voxels that will be retained in the final solution. The sparseness value can be determined empirically through cross-validations, or based on prior knowledge of the expected functional region size. Initially, we selected a fixed sparseness value of 0.045, which was determined from simulations with manually drawn ROIs at the beginning of the study. The manually drawn ROIs were simple spheres of approximately 3ml volume created in ITKsnap (Yushkevich et al., 2006).

Although simulations were first performed with a fixed sparseness value, this may not work for all situations. For example, a big functional unit might not be detected with our default sparseness value of 0.045. To adapt SCCAN for broader applications, we built an optimization routine that searches for the sparseness value with the best predictive accuracy. This is achieved by running 4 fold cross-validations (CV) at various sparseness values between 0.01 and 0.9. For each sparseness value, a model is built on 75% of the subjects and the behavioral scores are predicted for the remaining 25%. The sparseness value that achieves the highest CV correlation of predicted behavior with true behavior is selected for a final run of SCCAN with the full sample. This procedure is slower than a single SCCAN run but provides robust assessment of optimal sparseness. Moreover, it allows one to estimate the significance of the findings, i.e., a small CV correlation indicates that a reliable map cannot be built, and the result is null. Preliminary tests showed that sparseness could peak occasionally at high values, likely because the random split of the subjects required more voxels to reach the maximum CV correlation, or perhaps the random split of subjects was not optimal. Yet, smaller sparseness showed similar CV correlation. To enhance the accuracy of LSM maps, we added a penalty term to the optimization routine to penalize higher sparseness unless necessary. The penalty was set to 3% of the sparseness value. For example, the difference between sparseness 0.3 and 0.6 is 0.6*0.03 - 0.3*0.03 = 0.009. If the difference of CV correlations between these sparseness values was smaller than this value, the optimization would converge towards the smaller sparseness. It is important to note that sparseness does not change dramatically the pattern of results in itself. The effect of altering sparseness is visually similar to setting different cutoff thresholds on the final map. However, the optimization routine provides empirical evidence for the optimal extent of results.

### Measures of accuracy

We assessed the accuracy of the results with four measures. For each measure, the reference map was the binary mask of the brain parcel used for simulation, and the statistical map was compared with the reference map. The statistical map was binarized before the computations when necessary.

### Dice similarity (DICE)

This is a measure of spatial precision accuracy between 0 and 1, computed from binarized maps. The maximal value is achieved when the two maps overlap perfectly, and the number of false alarms (i.e., voxels outside the parcel of interest) is zero. A known limitation of dice overlap is its dependency on size, i.e., the same physical displacement leads to smaller dice in smaller brain regions but larger dice in larger brain regions (see (Pustina et al., 2016)).

### Average contour displacement (AC-D)

This is a metric of the average distance from the contour of the predicted map to the closest point of the brain parcel that generated the map. Different from dice, this measure is not sensitive to parcel size.

### Center of mass displacement (CM-D)

This measure has been used in previous studies to calculate the displacement of VLSM results (Mah et al., 2014; Sperber and Karnath, 2017). In our opinion, this measure is not optimal for measuring displacements because of the systematic bias towards the center of the analyzed area; i.e., a simulated cortical area can produce results only within the brain, producing centers of mass that are displaced only in one direction: toward subcortical areas. However, we computed CM-D for compatibility with previous studies. CM-D was computed as the euclidean distance between the center of mass of the binary brain region and the center of mass of the statistical map.

### Peak voxel displacement (PV-D)

LSM literature often focuses on the location of peak voxels. Thus, the displacement of the peak voxel has particular importance. We assigned PV-D a value of zero if the peak voxel fell within the simulated brain parcel, otherwise we computed the distance of the peak voxel to the closest point of the parcel that generated it.

### Statistical tests

After deriving accuracy measures for each of the 93 brain parcels, pairwise two-tailed Wilcoxon tests were performed for each accuracy measure to compare univariate vs. multivariate scores.

### Multi-region simulations

After confirming the value of SCCAN analyses for single brain regions, we performed multi-area simulations using the fully automated optimized SCCAN routine. To create multi-area simulations, six non-adjacent areas were selected in anterior, central, and posterior areas, both medially and laterally (areas 75, 96, 140, 144, 175, 2001; the last is a white matter tract). By combining these areas we created 15 two-area combinations and 15 three-area combinations. For each combination, behavioral simulations were created either with an “AND” rule, or with an “OR” rule. The AND rule consisted of averaging the lesion load of multiple regions before computing behavioral simulations. This is conceptually similar to having a large functional unit composed of separate regions, such that the damage of a region does not cause complete behavioral deficit unless the other regions are damaged as well. The OR rule consisted of computing the maximal lesion load for multiple regions and using this maximal lesion load to produce behavioral simulations. This is conceptually similar to interrupting a network of regions that are all crucial for performing a cognitive task. The OR rule corresponds to some extent to the “partial injury problem” explained in in Kinkingnéhun et al. (2007) and Rorden et al. (2009). All multi-area simulations were created with an error weight of 0.5.

Computations were performed on a server cluster operated under CentOS 6.6 with multiple Xeon E4-2450, 2.1GHz processors. The principal software used was R v3.1.1 with packages ANTsR (Avants, 2015), RcppArmadillo (Eddelbuettel and Sanderson, 2014), caret (Kuhn, 2008), lawstat (Hui et al., 2008), and nparcomp (Konietschke et al., 2015). Combined plots of accuracy measures were created with ggplot2 (Wickham, 2009).

## RESULTS

### Number of unique patches

The average number of patches at various resolutions are shown in Table 2. From these results it can be deduced that at every sample size there is a clear advantage of using high resolution lesion maps as opposed to lower resolution maps. The loss of unique patches can be seen at small samples sizes, with 2mm and 3mm maps containing respectively 67% and 48% of the unique patches of 1mm maps at N=20. The discrepancy becomes increasingly higher at larger sample sizes. At N=131 the number of unique patches obtained from 2mm and 3mm lesion maps is just 18% and 6% of patches that can be obtained with 1mm lesion maps.

**Table 2.** Number of unique patches at various resolutions and sample sizes. For samples smaller than the full sample (N=131) the cells contain the average number of patches in 200 randomly selected subsamples.

| Raw number of unique patches |  |  |  |  |  |  |  |  |  |  |  |  |
| --- | --- | --- | --- | --- | --- | --- | --- | --- | --- | --- | --- | --- |
|  | N20 | N30 | N40 | N50 | N60 | N70 | N80 | N90 | N100 | N110 | N120 | N131 |
| **1 mm** | 2803 | 11341 | 25301 | 43779 | 63986 | 86672 | 107825 | 129522 | 148560 | 166553 | 183146 | 195102 |
| **2 mm** | 1871 | 5932 | 10864 | 15947 | 20394 | 24484 | 27609 | 30417 | 32440 | 34146 | 35543 | 35709 |
| **3 mm** | 1358 | 3578 | 5671 | 7435 | 8729 | 9763 | 10433 | 11015 | 11359 | 11636 | 11850 | 11602 |
| Percentages |  |  |  |  |  |  |  |  |  |  |  |  |
|  | N20 | N30 | N40 | N50 | N60 | N70 | N80 | N90 | N100 | N110 | N120 | N131 |
| **1 mm** | 100% | 100% | 100% | 100% | 100% | 100% | 100% | 100% | 100% | 100% | 100% | 100% |
| **2 mm** | 67% | 52% | 43% | 36% | 32% | 28% | 26% | 23% | 22% | 21% | 19% | 18% |
| **3 mm** | 48% | 32% | 22% | 17% | 14% | 11% | 10% | 9% | 8% | 7% | 6% | 6% |

### Univariate vs. multivariate analyses

Highly localized findings were produced with SCCAN (fixed sparseness = 0.045), which were proximal or inside the simulated region, whereas BM produced statistical maps that extended frequently beyond the simulated area. Figure 2 displays statistical maps for four representative brain regions using SCCAN and BM with various thresholding mechanisms. A complete <u>video with all 93</u> regions is available as Supplementary Material. Overall, the most accurate univariate results were obtained with FWER multiple comparison correction. However, even these maps were not as precise and accurate as SCCAN maps.

**Figure 2.**
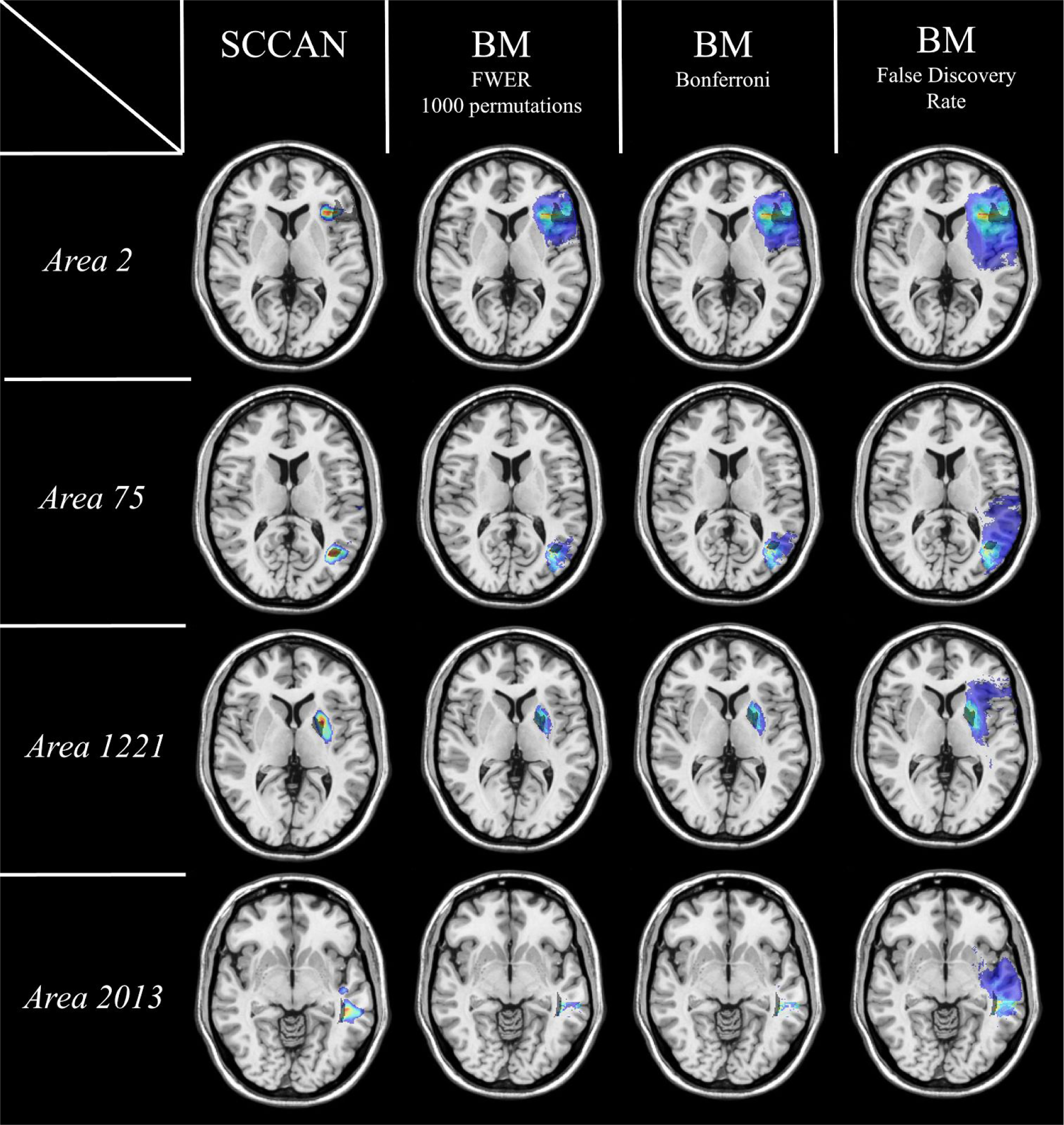
Representative LSM maps obtained from 4 brain regions with SCCAN (1st column) and Brunner-Munzel tests at various thresholds (columns 2-4). All univariate analyses were thresholded at p < 0.05. Dark semi-transparent voxels show the true location of the simulated region.

Figure 3 displays plots comparing SCCAN and BM methods for all 93 simulated regions. The systematic advantage of SCCAN is clearly evident from these plots. Statistical comparisons showed higher dice overlap and less average displacement with SCCAN compared to BM maps thresholded with FDR, Bonferroni, or FWER (all p < 0.001). Compared to SCCAN, the displacement of the center of mass was higher for BM corrected with FDR or Bonferroni (p = 0.001 and p < 0.003, respectively), but not different for BM corrected with FWER (p = 0.13). Peak voxel displacement was the only measure that showed no difference between the various analyses and correction mechanisms (all p > 0.1). Overall, these results show that SCCAN is more accurate than the mass-univariate approach despite thresholding univariate results with various thresholding mechanisms.

**Figure 3.**
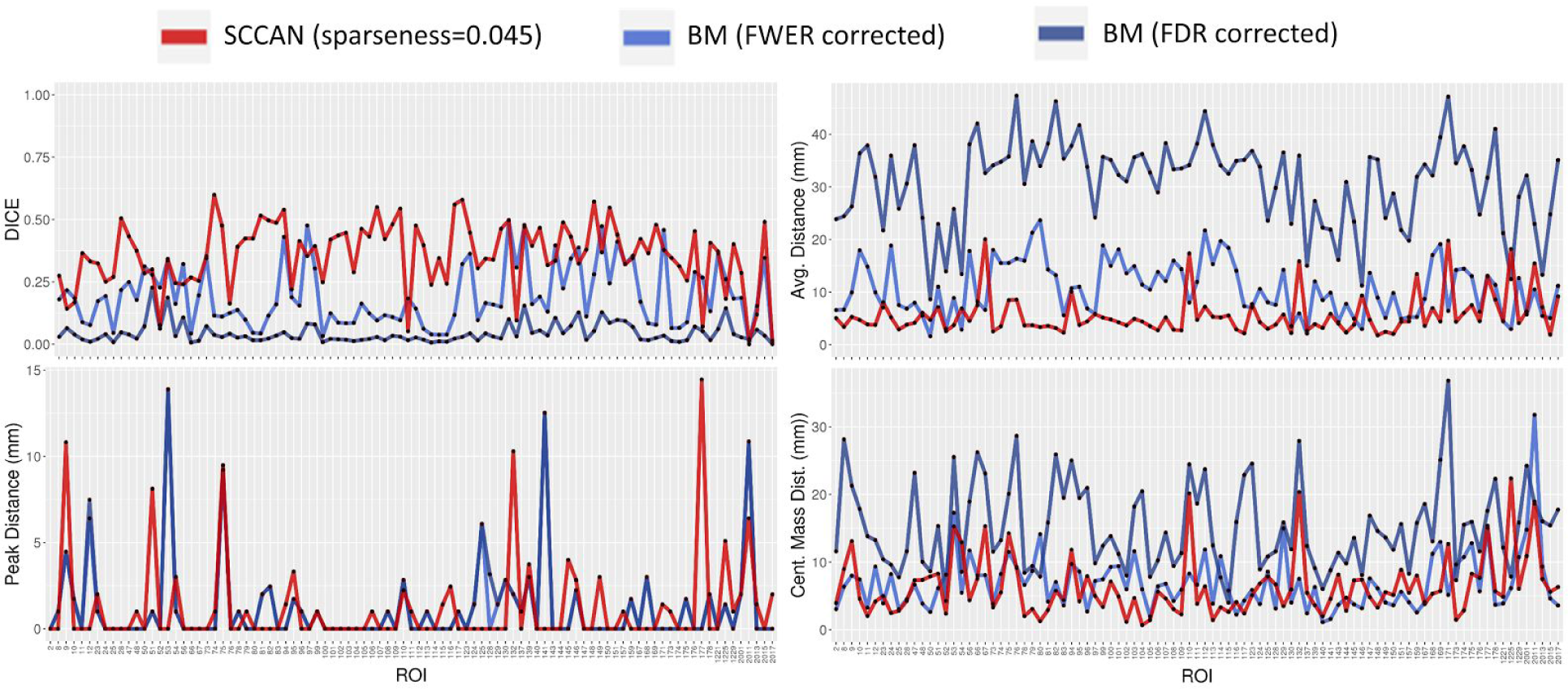
Line plots of the four accuracy measures for SCCAN (red) and BM (blue) methods.

The next question we asked was how would univariate and multivariate approaches compare at different sample sizes. To answer this question, we ran tests with reduced samples sizes of N = 20, 40, 60, 80, and 100. For each sample sizes, a single random subset of subjects was selected from the full dataset, and analyses were performed using the previously simulated behavioral scores. As before, only voxels lesioned in at least 10% of subjects were analyzed, and only brain regions affected in at least 10% of subjects with at least 50% damage in one subject were tested. Figure 4 displays dice scores of SCCAN and BM for all regions. Statistical comparisons were performed between SCCAN and BM (FDR or FWER thresholded) at every sample size. Dice overlap was significantly higher for SCCAN for all sample sizes (all p < 0.001). Starting at N=40, SCCAN showed a clearer advantage over BM all the way up to N=100. Typically, this was reflected in higher dice, lower average displacement, and lower center of mass displacement. The two methods, however, were frequently similar regarding the displacement of the peak voxel.

**Figure 4.**
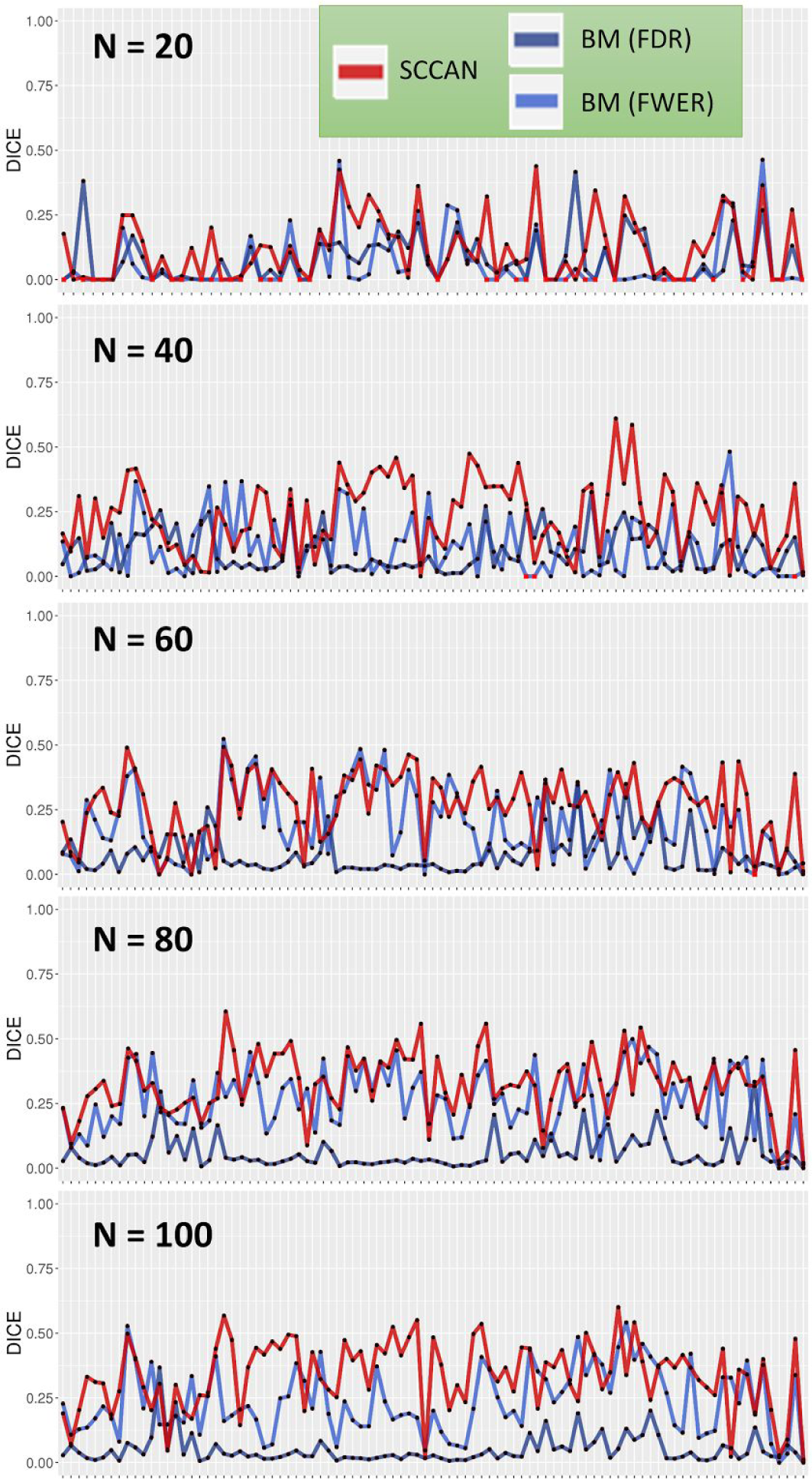
Dice overlaps of SCCAN (red) and BM (FDR thresholded: dark blue, FWER thresholded: light blue). The number of regions can be different between sample sizes. A red dot represents an analysis that produced no results (i.e., empty map). All statistical comparisons showed that SCCAN has higher dice score at every sample size (all p < 0.001).

### Fully optimized SCCAN sparseness

Before running multi-area simulations, we tested the efficiency of the optimization routine on the 93 single-area simulations (N=131). These analyses produced an average CV correlation of r = 0.72 +/- 0.1 (range: 0.40 - 0.85, median: 0.74) and selected sparseness values around 0.063 +/- 0.050 (range: 0.024 - 0.236, median: 0.036). Note that these sparseness values are close to the fixed value we used for the initial simulations. The maps produced with fully automated SCCAN had better dice overlap and average displacement than BM FWER thresholded maps (all p < 0.001). The advantage was not found in center of mass and peak displacement (p > 0.1). A comparison of fully automated SCCAN with our previous SCCAN results at fixed sparseness showed a small but significant gain in dice overlap for the automated method (p = 0.018). The other accuracy measures showed no difference between SCCAN runs.

### Multi-area simulations

The results from multi-area simulations cannot be properly measured with the four accuracy measures. For example, results that find one simulated area and miss all the others can yield a small average displacement because of the proximity with the identified area. The concept of peak distance is also not useful in this scenario because there are multiple areas under investigation. For this reason we considered only dice overlap for comparisons between methods. Figure 5 displays the resulting plots for two- and three-area simulations. When compared with BM FDR corrected, SCCAN was always more accurate (all p < 0.001). When compared with BM FWER corrected, SCCAN was more accurate only for three-area combinations with an AND rule (p < 0.001). We noticed, however, that SCCAN could detect more frequently the simulated regions, while BM failed find to find simulated regions a larger number of times. For three-area AND combinations BM failed to find almost half of the regions; i.e., 21 missed regions out of 45 simulated regions. For other combinations a similar tendency was observed, though individual chi-square tests were not significant. These results show that the two methods produced two different patterns, one which found the areas but showed low dice overlap because of larger maps (SCCAN), the other which showed limited results and missed the simulated regions more frequently (BM-FWER). Note that, as opposed to BM, SCCAN findings are based on cross-validated optimization. Example maps showing results from SCCAN and BM are displayed in Figure 6.

**Figure 5.**
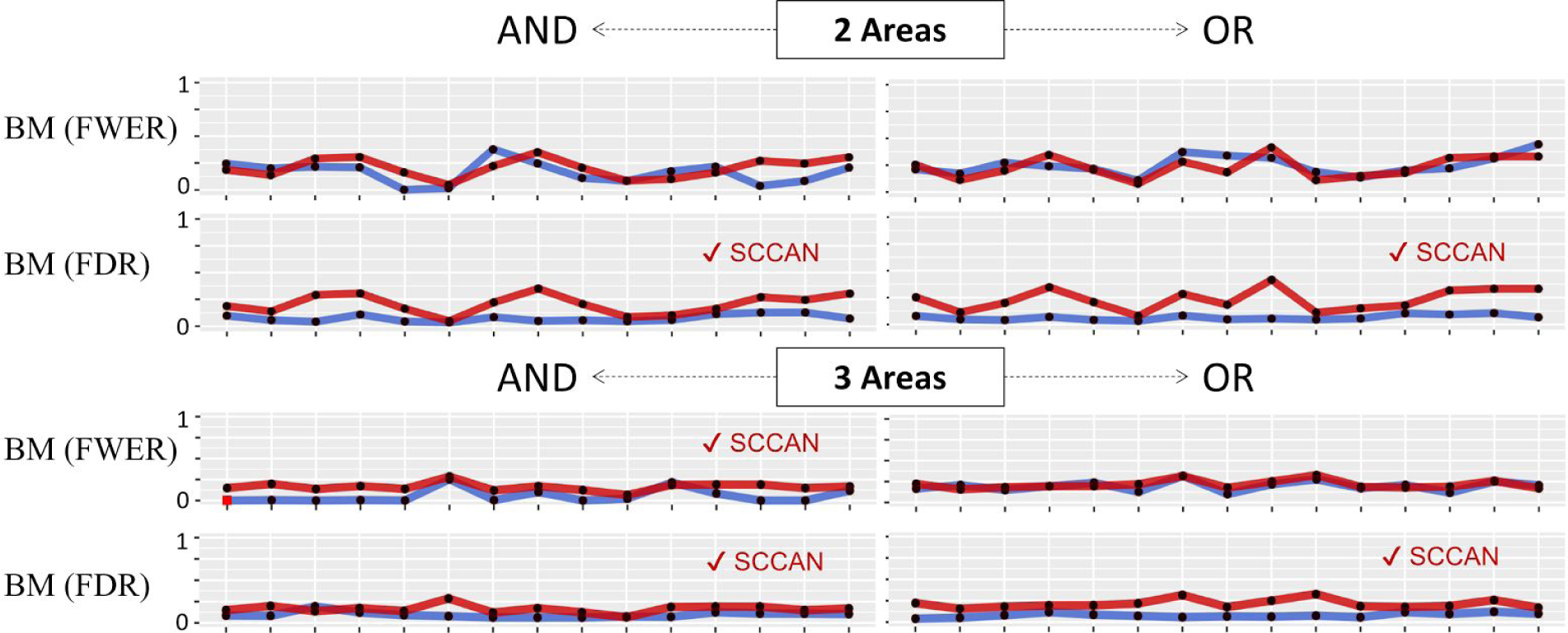
Dice overlap of results from SCCAN (red) and BM (blue) analyses. Checkmarks on the side indicate which method was statistically superior (all p < 0.002). A red dot in the plot indicates that the analysis failed to produce any result.

**Figure 6.**
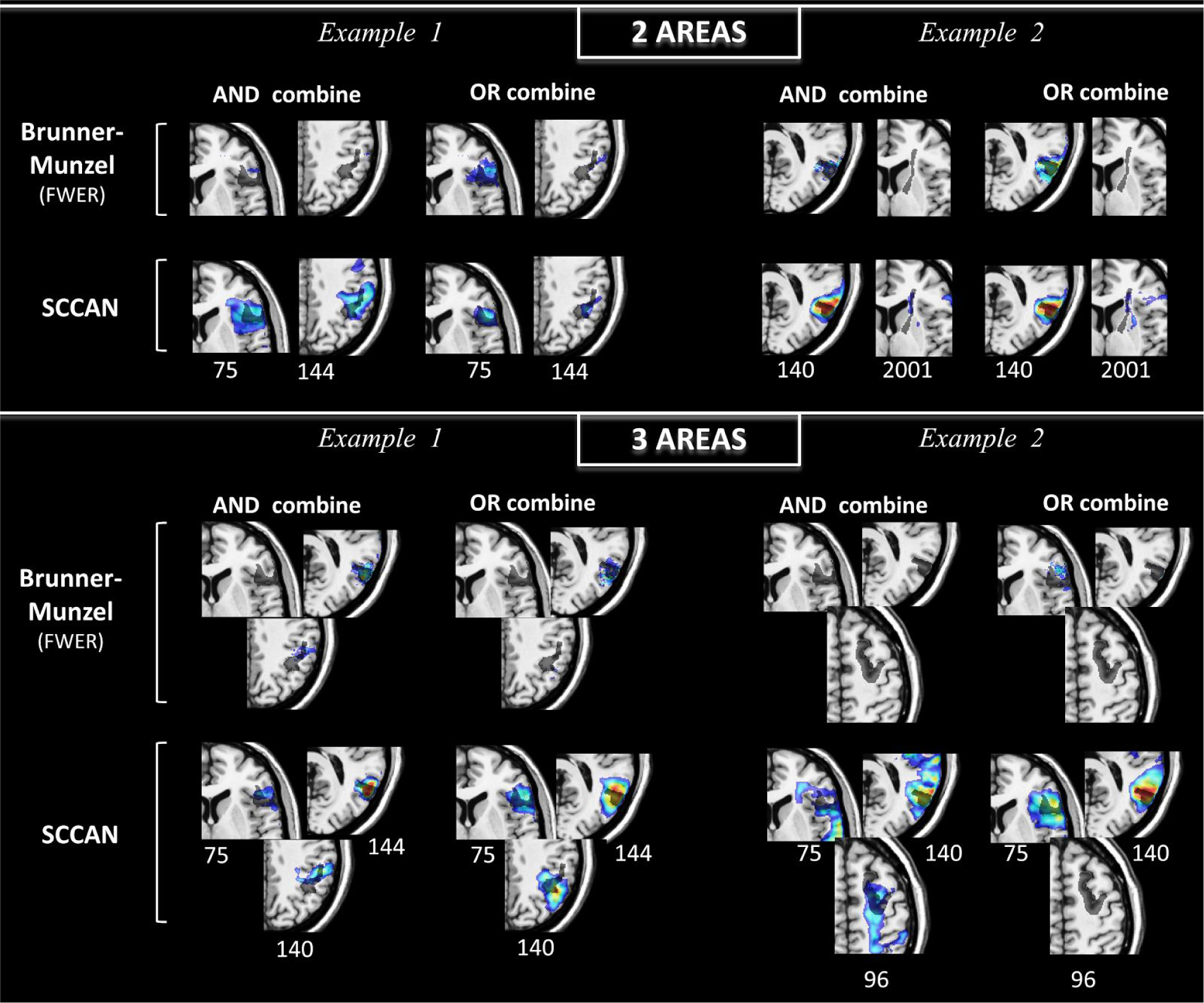
Example results from two-area (above) and three-area (below) combinations. The range of the colormap extends at full extent for each analysis. The numbers below the maps indicate the parcel number. The parcel itself is displayed in dark semi-transparent color.

Descriptive statistics of multi-area results are listed in Table 3. The cross-validated correlation from multi-area SCCAN was always above the statistical threshold (all p < 0.001).

**Table 3.**
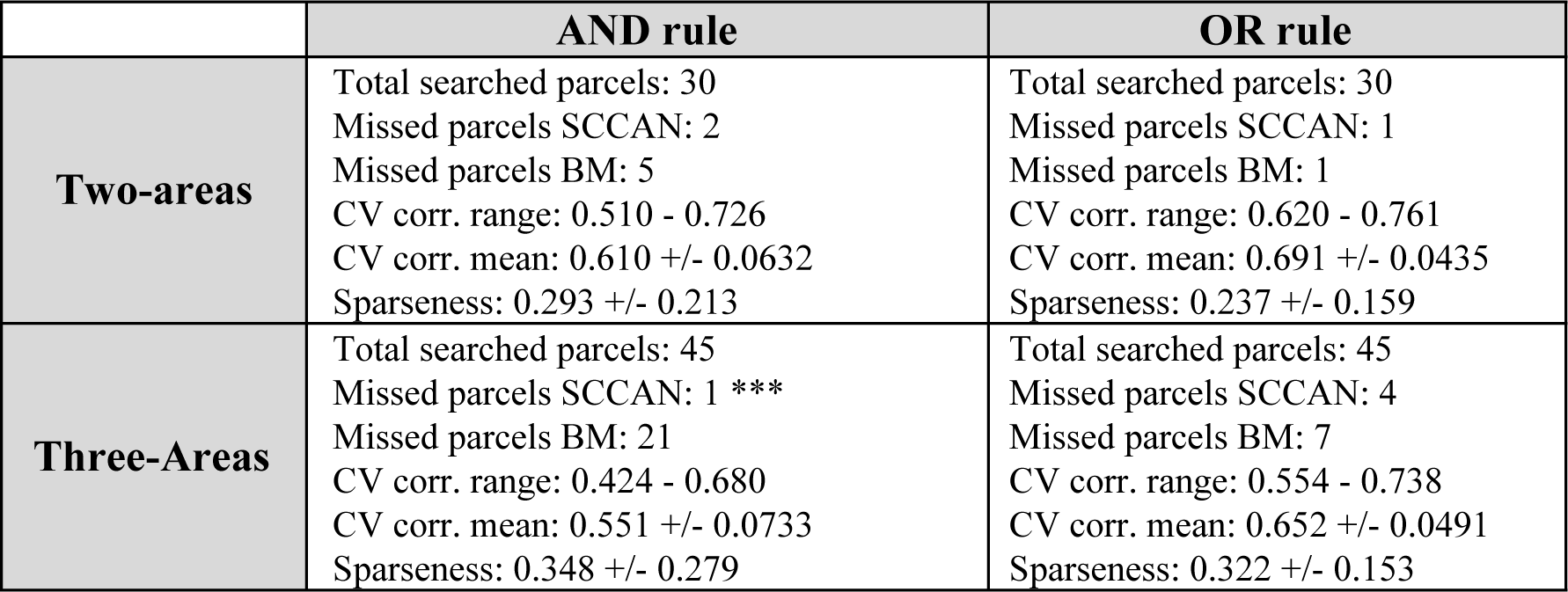
Descriptive statistics of the cross-validated correlation for the multi-area combinations obtained with SCCAN. Missed parcels show the number of the simulated parcels that had no significant voxels, and, therefore, were completely missed. CV corr = cross-validated correlation, *** (three asterisks) denote a significant difference between SCCAN and BM-FWER for the number of missed parcels (chi square test, p<0.001).

|  | AND rule | OR rule |
| --- | --- | --- |
| **Two-areas** | Total searched parcels: 30<br>Missed parcels SCCAN: 2<br>Missed parcels BM: 5<br>CV corr. range: 0.510 - 0.726<br>CV corr. mean: 0.610 +/- 0.0632<br>Sparseness: 0.293 +/- 0.213 | Total searched parcels: 30<br>Missed parcels SCCAN: 1<br>Missed parcels BM: 1<br>CV corr. range: 0.620 - 0.761<br>CV corr. mean: 0.691 +/- 0.0435<br>Sparseness: 0.237 +/- 0.159 |
| **Three-Areas** | Total searched parcels: 45<br>Missed parcels SCCAN: 1 ***<br>Missed parcels BM: 21<br>CV corr. range: 0.424 - 0.680<br>CV corr. mean: 0.551 +/- 0.0733<br>Sparseness: 0.348 +/- 0.279 | Total searched parcels: 45<br>Missed parcels SCCAN: 4<br>Missed parcels BM: 7<br>CV corr. range: 0.554 - 0.738<br>CV corr. mean: 0.652 +/- 0.0491<br>Sparseness: 0.322 +/- 0.153 |

### Avoiding spurious results

One of the most important aspects of any method is the ability to avoid false positive findings. SCCAN showed good accuracy when presented with a verified brain behavior relationship. But how would it perform when there is no relationship in the data? To test this scenario, we ran SCCAN after scrambling one of the simulated behavioral scores. After searching for the optimal sparseness value, the algorithm found poor cross-validated correlation (r=-0.07, p=0.4), and returned a null finding (empty statistical map). This test showed that SCCAN could correctly reject a randomly created brain behavior relationship.

### Analyzing real behavioral scores

After completing all simulations, we analyzed two real scores, WAB-AQ and PNTcorrect. Figure 7 displays sagittal slices showing univariate and multivariate results. Overall, SCCAN identified specific regions, while BM tests either produced diffuse maps (FDR thresholded) or limited results (FWER thresholded, 1000 permutations). The cross-validated correlation of SCCAN was 0.55 and 0.42 for WAB-AQ and PNTcorrect, respectively. For WAB-AQ, SCCAN identified two major areas related to the deficit: the rolandic operculum near the parietotemporal junction (i.e., area Spt, (Hickok et al., 2003; Schwartz et al., 2012)), and the subcortical frontal white matter (Dronkers et al., 2007; Fridriksson et al., 2013). For picture naming, the pattern of findings were similar, but the findings extended further to include anterior temporal and parietal regions (i.e., angular gyrus). A subtraction of PNT and WAB maps exposed more clearly the latter two areas as being more strongly related to picture naming than to the aphasia quotient.

**Figure 7.**
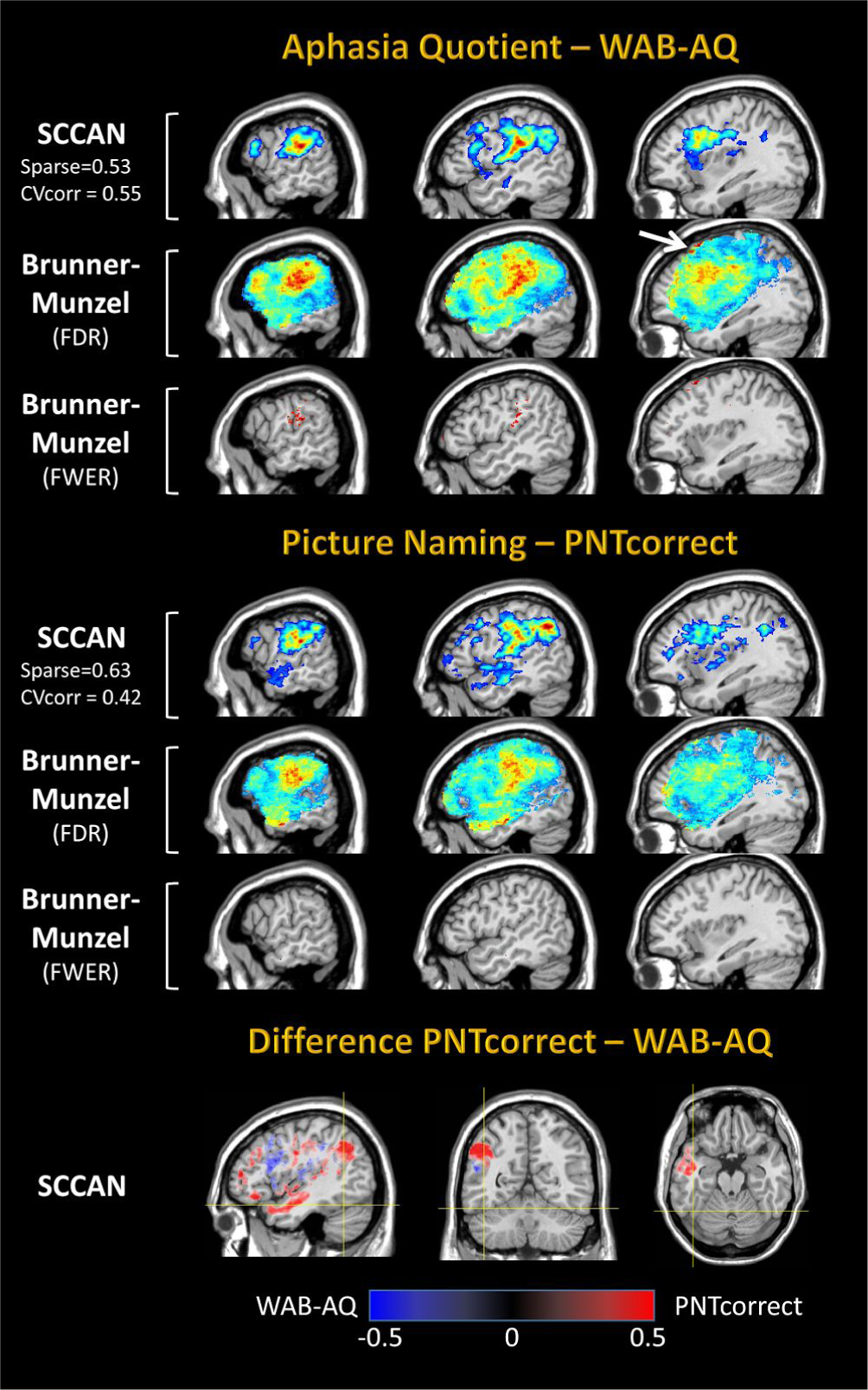
LSM analyses of two behavioral scores, WAB-AQ (above) and PNTcorrect (below) with univariate and multivariate methods. Colormaps extend to the full range of the statistical findings for each method.

## DISCUSSION

We introduced a novel multivariate method for lesion to symptom mapping (SCCAN) and compared it with traditional state of the art mass-univariate analyses. A wide spectrum of potential brain-behavioral relationships were simulated, including functional dependency on single and multiple areas, different sample sizes (N=20-131), and different regional combinations (i.e., AND, OR). We also assessed more than one univariate thresholding mechanism (FWER, Bonferroni, FDR) and compared the methods with real behavioral scores from aphasic patients. Overall, SCCAN demonstrated improved spatial accuracy compared to univariate tests for almost all the comparisons.

The SCCAN method was born out of traditional canonical correlation formulations (Hotelling, 1936) and was recently extended for use with neuroimaging data (Avants et al., 2010). SCCAN has been successfully used to identify the complex relationships between brain properties and cognitive scores in patients with neurologic diseases (Avants et al., 2014, 2010; McMillan et al., 2014), and to relate structural and functional brain properties (Avants, 2017). In this work, we extended the use of this technique to lesion-to-symptom mapping, and created optimization procedures to enable it for routine use.

The quest of multivariate LSM follows an increased acknowledgement of the limitations of the mass-univariate approach (Karnath and Smith, 2014; Kimberg et al., 2007; Mah et al., 2014; Nachev, 2015; Sperber and Karnath, 2017). For example, the variation of statistical power across the brain in lesion studies can cause greater difficulty to detect functional units located in areas lesioned infrequently compared to functional units located in areas lesioned frequently^1^ (Kimberg et al., 2007; Rudrauf et al., 2008). Such variation in statistical power may interact with the natural variation of effect size to create complex scenarios; i.e., a weaker effect size can be detected more promptly when located in an area with high statistical power. Even if statistical power was homogeneous, the high correlations between neighboring voxels create diffuse statistical maps with limited spatial accuracy. Confirming these concerns, we found that the average correlation of a lesioned voxel with its immediate neighbors is r = 0.88 +/- 0.05. When faced with diffuse results, the only choice researchers have available is the variation of statistical threshold such that less (or more) voxels are deemed significant. However, no thresholding approach can handle all scenarios, nor can voxel-wise mass univariate methods capture distributed (network-like) relationships. For example, in our simulations FWER correction was the most accurate thresholding mechanism, but its application to real behavioral scores produced almost non-existent findings. This lack of results cannot be readily interpreted because all patients had some degree of aphasia caused by stroke, and, therefore, a relationship between aphasia severity and lesion location should have emerged. Even out of this practical scenario, FWER thresholding makes the implicit assumption that voxels are exchangeable (Nichols and Holmes, 2002; Winkler et al., 2014). Yet, the variation in statistical power makes this assumption more difficult to satisfy.

The limitations of mass-univariate VLSM have been shown in several recent studies. For example, Mah et al. (2014) showed that the center of mass of activation maps is frequently shifted towards subcortical areas where lesions are more frequent and the statistical power is higher. Sperber and Karnath (2017) confirmed these results and identified factors that partially mitigate the problem, such as, using voxels lesioned in more than 5% of subjects, or regressing out the effect of lesion size. While the above simulations rely mostly on simulations of single brain regions (or single voxels), higher cognitive skills typically involve a network of structures (Medaglia et al., 2015; Mill et al., 2017; Petersen and Sporns, 2015), and, therefore, multi-area simulations are very important. In this regard, Mah et al. (2014) showed that univariate analyses may fail to find functional units located in multiple brain regions. Zhang et al. (2014) performed three-area simulations and found that univariate methods can miss some of the simulated regions. At a conceptual level, multivariate methods are better equipped to overcome the above limitations. The simple explanation is that, since all voxels are considered together in a single model, there are better chances of achieving correct *mapping* of the voxels that participate in behavioral performance. This benefit has been shown both in Mah et al. (2014) and in Zhang et al (2014). Specifically, these authors have shown that multivariate analyses based on support vector regression (SVR) can outperform traditional VLSM in detecting multi-area combinations. In Mah et al, this effect was deduced from a real analysis, while in Zhang et al. the results were obtained from simulated cubes of the same size with perfect brain-behavior relationships (i.e., no noise in the simulations). The multivariate advantage found by Zhang et al. appeared only after correcting the data for lesion size, but not when raw data were used for analyses. In our simulations, we never corrected for lesion size, and despite this SCCAN outperformed VLSM. Thus, our results extend prior conclusions by showing that another multivariate solution, SCCAN, can outperform univariate analyses in the context of one-area, two-area, and three-area, noisy simulations.

The creation of different types of multi-area simulations (“AND”, “OR”) was a novel approach with particular importance for future LSM studies. The “AND” and “OR” combinations cover two basic principles of interactions between brain regions within a functional network. The “AND” rule assumes that lesion of an area is insufficient to produce behavioral deficits because functional compensation may occur in another area. Conceptually, regions can be considered a functional extension of each other, just like having a larger functional area which is composed of non-adjacent voxels. The process of functional compensation is thought to be one of the main mechanisms of cognitive recovery after stroke, and, consequently, the “AND” combination is potentially important for considering intervention and recovery mechanisms (R. H. Hamilton et al., 2011; Hope et al., 2017; Piai et al., 2017; Tracy et al., 2014; Turkeltaub et al., 2012). From an information flow perspective, “AND” combinations can be considered as parallel pathways of information processing into different areas. The simulations performed by Zhang et al. (2014) were similar to the “AND” combinations we used here. On a different plane, the “OR” rule considers all functional areas as critical nodes in a network, such that the lesion of an area would shut down the entire processing ability. From an information flow perspective, the “OR” combination is equivalent to a serial processing stream (Dell et al., 1997; Schwartz et al., 2006/2). The “OR” rule can also be thought of as the “partial injury problem” previously described in Kinkingnéhun et al. (2007) and Rorden et al. (2009). Altogether, the “AND” and “OR” combinations provide good coverage of the four mechanisms of single- and multi-area brain-behavior relationship described in Godefrey et al. (1998) (unicity, equivalence, association, summation). SCCAN was sensitive at detecting both “AND” and “OR” combinations, albeit occasionally some parcels were missed. Typically more parcels were missed with BM analyses (Table 3). This difference was particularly clear with the three-area “AND” combinations, for which almost half of all the simulated brain regions were missed by the univariate method.

Univariate analyses were, however, robust with “OR” combinations, ultimately producing similar accuracy as SCCAN. These findings suggest that “AND” relationships between brain regions are more at risk of going undetected by traditional VLSM than “OR” relationships. From an information flow perspective, this is equivalent to having poorer detection of parallel pathways compared to serial pathways. Overall, there was no combination in which the univariate method would be preferable over SCCAN, the multivariate method would either outperform univariate analyses or perform equally well.

Among the accuracy measures used to compare the methods, the peak voxel location was the most resilient to method limitations, and was frequently similar between SCCAN and BM analyses. This is good news because the peak voxel is frequently used to find the brain area maximally related to the behavioral score. Yet, the peak voxel can also be misleading, not only because it can be displaced due to variations in power and effect size, but also because previous evidence has shown that variations in lesions maps can cause occasionally large displacements of peak values (Pustina et al., 2016). Moreover, the value of the peak voxel is useful only when a single brain region is the source of the behavioral deficit. Proper mapping of multiple broader brain regions require methods that provide spatial accuracy independently of where the peak voxel is located.

The SCCAN analyses of true behavioral scores (i.e., aphasia quotient and picture naming) revealed several areas implicated in language deficits. Although a detailed interpretation is beyond the scope of this paper, it is worth noting that SCCAN mapped behavioral deficits to lesions located in known language processing areas. For example, aphasia severity was related to damage of the parieto-temporal junction and the frontal white matter. The parieto-temporal junction is an area involved in phonological retrieval or auditory motor interaction (Hickok et al., 2003), while frontal white matter has been frequently found as a crucial bottleneck whose lesion causes multiple aphasia symptoms (Dronkers et al., 2007; Fridriksson et al., 2013; Griffis et al., 2017; Ivanova et al., 2016). Picture naming showed additional involvement of lesions in anterior temporal and parietal areas (Figure 7). The anterior temporal lobe is a known hub for semantic processing, as evidenced from lesion studies (Fridriksson et al., 2016; Halai et al., 2017; Harvey and Schnur, 2015; Schwartz et al., 2009), other neurologic diseases (e.g., epilepsy (Lambon Ralph et al., 2012), and healthy controls (Geranmayeh et al., 2015). Thus, the difference between WAB-AQ and PNTcorrect maps we discovered might be related to the stronger semantic component of the picture naming task.

Our investigation of the number of unique patches suggested that high resolution maps can provide more detailed information than maps downsampled at 2mm or 3mm. Note that having more unique patches does not necessarily resolve the problem of spatial correlation between neighboring voxels. Neighboring voxels can still have high correlation values. However, the diversity in the data might be advantageous to multivariate methods. The ultimate conclusion whether voxel diversity is useful or it just adds an unjustified number of multiple comparisons requires additional simulations that can be performed in future studies. This question is particularly relevant for mass-univariate analyses adopted in current VLSM practices, but may also deserve simulations in a multivariate context.

This study was an initial attempt to test the value of SCCAN for lesion to symptom mapping without performing an exhaustive search on all the parameters. Future work is needed to elucidate better the effects of smoothness, number of iterations, gradient descent parameter, orthogonalization, and rank transformations. Some of these parameters may further improve the accuracy while others may lead to faster processing. Understanding the relationship between sparseness values and spatial accuracy may allow the creation of sparseness curves that describe the link between localization precision and loss of predictive power, ultimately finding a better penalty value (currently 0.03). Another aspect which deserves more attention is the introduction of non-linearities in the simulations. Evidence from previous literature shows that true relationships between lesion load and behavioral deficit might have a strong non linear component (Wang et al., 2013). We have enabled the simulation tool in LESYMAP to produce non-linear relationships, but did not aim to investigate this aspect in the present work. A natural extension of this work is the comparison of SCCAN with other multivariate methods, such as SVR. Although we did not compare these methods, future tests will compare both of these methods.

Besides developing multivariate methods, efforts directed at improving existing univariate methods might have merit. For example, neighborhood correlation maps can be used to characterize differences between brain regions (Rudrauf et al., 2008). One can also envision attempts to implicitly correct statistical maps for the variation in power, or other models of correction similar to the one proposed by Sperber and Karnath (2017).

The tools utilized in this study are available in the LESYMAP package for R. LESYMAP relies on ANTsR (Avants, 2015) for image manipulation and SCCAN analyses, but contains a number of custom functions for running traditional VLSM analyses (t-test, regression, BM). All thresholding methods mentioned in this study are available in LESYMAP. Within this package, C++ functions allow fast permutation-based thresholding methods (i.e., FWER). An FWER routine with 1000 permutations and 200,000 voxels takes 20-60 minutes, depending on computer architecture. SCCAN is typically slower to run, and its speed depends on several factors. Generally, a single SCCAN run takes 5-15 minutes, while a complete SCCAN run with optimization takes 2-12 hours. Typically, smaller functional regions require smaller sparseness to be detected, and this translates to more time. A complete SCCAN run provides not just a map, but also a measure of cross-validated correlation. This measure can be thought of as an overall measure of the strength of the findings and represents the confidence of the model. If behavioral deficits arise from non-corresponding areas in different patients, the CV correlation will be low, indicating that the result might be difficult to replicate in other samples. In this respect, CV correlation represents a departure from classical voxel based thresholds in favor of a full map indicator of predictive accuracy. We believe this approach will enhance the reproducibility of future LSM studies.

Although we controlled a number of factors, there are several limitations worthy of note. We investigated only a few of the possible factors that influence LSM results. Future work may focus on additional investigations of non-linearity, sample sizes, functional unit sizes, etc. Ultimately, the number of factors to investigate is large and difficult to cover in a single study. Second, CV correlation is used only to select the sparseness value, not to verify the predictive power of the final map. The predictive value of sparseness indicates that there is a certain brain pattern obtained with a specific sparseness that is predictive of the behavior of other subjects. Yet, that brain pattern may vary slightly at each cross-validation run. Strictly speaking, the final map does not guarantee the same exact predictive power obtained when optimizing sparseness.

In this regard, the aim of the current software version (LESYMAP v0.0.0.9001) is to offer a lesion-to-symptom mapping tool, and not a predictive tool. Third, a risk inherent to newly developed tools is the possibility of bugs and errors. We performed several checks and comparisons with existing software (MRIcron and Voxbo, (Kimberg et al., 2007; Rorden et al., 2007)), and repeated the analyses when necessary, performing over 16,000 LSM analyses throughout the study. Yet, as history have shown (Medina et al., 2010) there might be bugs in the software that will be discovered later. Finally, it should be noted that some of the results are based on a single random selection of subjects from the full sample. Normally this random selection should be repeated many times to avoid sample selection bias, but the computational costs of these repetitions are high. Thus, some of our findings may contain random error from subject selection (i.e., another set of 20 subjects might have produced slightly different results when we tested LSM at N=20).

In conclusion, we started this study by asking whether multivariate methods for lesion-to-symptom mapping can produce more accurate results, as some authors had suggested (Karnath and Smith, 2014; Mah et al., 2014; Nachev, 2015; Zhang et al., 2014). We found that this was indeed the case, and that SCCAN can outperform traditional VLSM analyses in most of the tested scenarios. The public availability of this method will allow researchers to collectively draw firmer conclusions on the spatial topography of cognitive systems. At the same time, SCCAN opens new possibilities for investigating functional organization theories by identifying entire cognitive components derived from all behavioral scores (Fridriksson et al., 2016; Mirman et al., 2015). In this way, SCCAN represents a harmonization between the statistical theory behind construct psychology and multivariate neuroimaging analysis, which defines a frontier of modern cognitive neuroscience. Researchers can extend the functionality of LESYMAP with other LSM methods such as support vector regression or random forests (i.e., see the advantage of decision trees in (Godefroy et al., 1998)). This work is a small step towards more accurate and reproducible lesions studies. The LESYMAP package is available online at https://github.com/dorianps/LESYMAP.

## Footnotes

1 Technically the power goes whether a voxel has too few or too many lesions, but voxels are seldom lesioned in more than 50% of the subjects.

## ACKNOWLEDGEMENTS

We would like to thank Myrna Schwartz for initial comments on the manuscript, Laurel Buxbaum and Erica Middleton for agreeing to share the lesion maps with the public, Dan Mirman for testing LESYMAP, and Pati Sarthak for assisting with C++ functions.

---

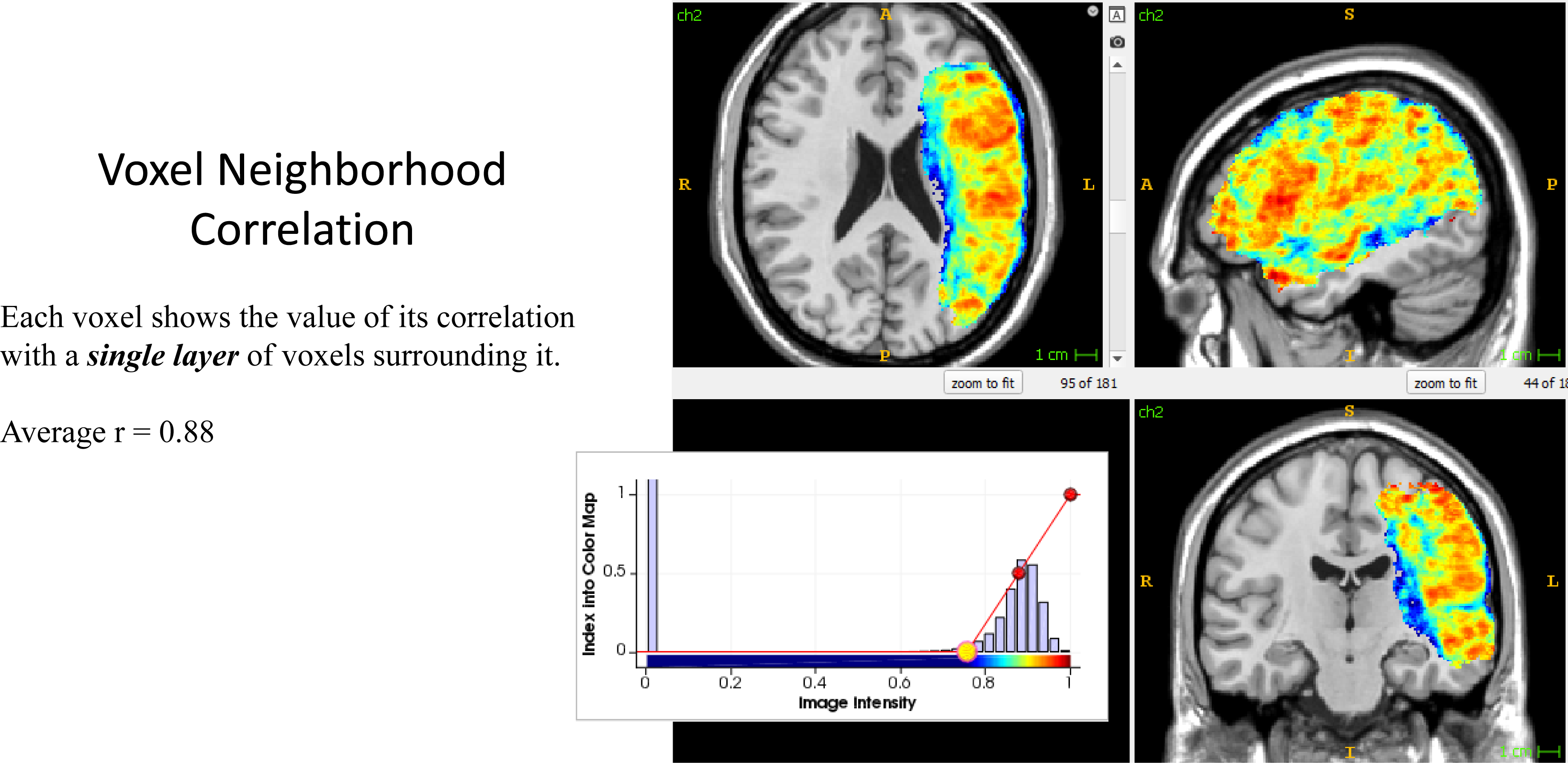

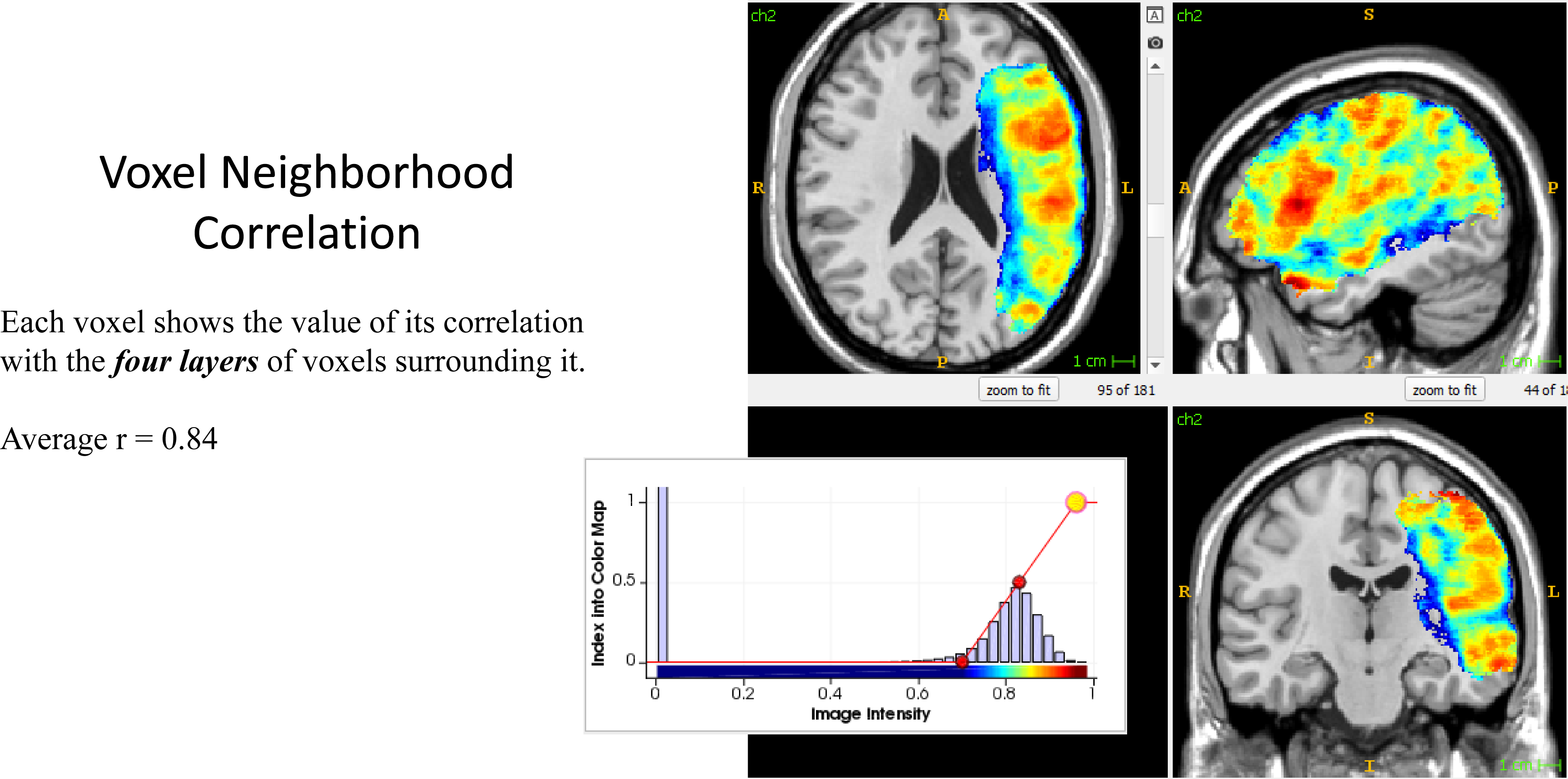

